# Native Mass Spectrometry Reveals Distinct Effects of Linker Length and Targeting-Ligand Identity on PROTAC Ternary Complexes

**DOI:** 10.64898/2026.09.29.755516

**Authors:** Evan N. Whitford, Joshua D. Gilbert, Marius M. Kostelic, Michelle A. Estrada, Aaron C. Ehlinger, Tiffany A. Thibaudeau, Shaun M. McLoughlin, Vicki H. Wysocki

## Abstract

Proteolysis-targeting chimeras (PROTACs) promote targeted protein degradation by inducing a transient ternary complex between a protein of interest and an E3 ubiquitin ligase. Despite the clinical promise of these event-driven therapeutics, defining the conformational landscape and interfacial organization of these complexes remains challenging. Here, we use an integrated native mass spectrometry (nMS) approach combining ion mobility, surface-induced dissociation (SID), and variable-temperature electrospray ionization (vT-ESI) to determine how small-molecule architecture shapes the structural and thermodynamic landscapes of bromodomain-containing protein 4 (BRD4) with the VCB E3 ligase complex. Using a set of PROTACs that differ in targeting ligand identity (JQ1 versus I-BET726) and PEG linker length, ion mobility measurements show that shorter linkers constrain the ternary complex into more extended topologies, whereas longer, more flexible linkers allow dynamic sampling of relaxed, native-like conformations. SID directly probes noncovalent interfaces and shows that JQ1 complexes favor prompt PROTAC ejection, whereas tighter-binding I-BET726 complexes are more likely to retain the PROTAC at the interface. Complementing these kinetically-controlled gas-phase measurements, vT-ESI reveals a monophasic melting transition for a JQ1-mediated complex and a distinct biphasic melting profile for an I-BET726-mediated complex. The gas-phase activation measurements revealed targeting-ligand-driven differences in PROTAC ejection and retention, whereas vT-ESI uncovered distinct temperature-dependent disassembly pathways for the matched-linker-length JQ10L and IBET10L complexes. Together, these complementary techniques show how PROTAC architecture influences ternary-complex conformation, interfacial connectivity, and disassembly.

**Synopsis:** Proteolysis-targeting chimeras (PROTACs) offer a promising therapeutic strategy by selectively directing disease-related proteins for intracellular degradation. Here, we employ a systematic native mass spectrometry approach to investigate how PROTAC design impacts the conformation and stability of the resulting ternary complexes. Integrating gas- and solution-phase data provides critical insights to inform rational PROTAC optimization.

## Introduction

Proteolysis-targeting chimeras (PROTACs) are a class of heterobifunctional small molecules designed to induce targeted protein degradation, offering a powerful strategy in drug development. Their design, which includes a targeting ligand for the protein of interest (POI), a targeting ligand for the E3 ligase, and a chemical linker connecting the ligands, enables the selective degradation of proteins previously considered undruggable, thereby broadening the therapeutic landscape and offering the potential for more durable and effective clinical outcomes.^1–4^ Unlike conventional inhibitors that rely on occupying active sites, PROTACs operate via an event-driven mechanism in which they simultaneously engage a therapeutic POI and an E3 ubiquitin ligase, thereby bringing them into proximity.^5,6^ This ternary complex formation promotes ubiquitination of the POI, marking it for recognition and subsequent degradation by the endogenous ubiquitin-proteasome system (Figure 1).^7^ By exploiting the cell’s endogenous degradation machinery, PROTACs can target proteins that lack accessible binding pockets, remain effective against some target-site mutations that confer resistance to conventional inhibitors, and produce sustained effects at lower drug exposure.^4,8,9^

**Figure 1:**
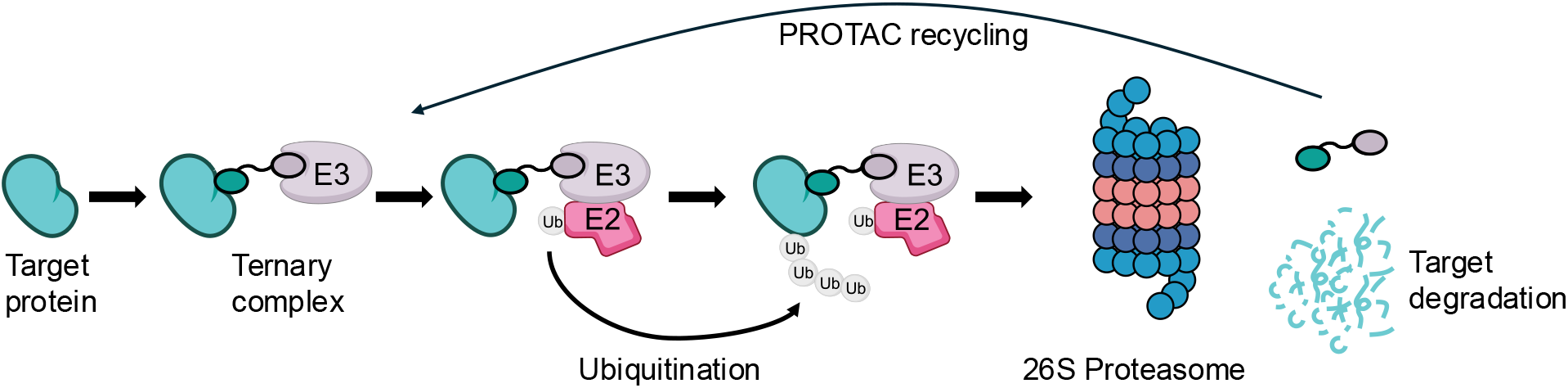
Scheme of PROTAC-induced targeted protein degradation. PROTACs induce a ternary complex between target protein and E3 ligase complex. The ternary complex facilitates ubiquitin conjugation to the target protein, enabling polyubiquitination and proteasomal degradation of the target protein.

The von Hippel-Lindau (V) protein, in complex with elongin B (eloB, B) and elongin C (eloC, C) (collectively referred to as VCB), is a widely employed E3 ligase in PROTAC design.^10^ Upon binding to either the ligase or the target protein of interest, a PROTAC forms a binary complex that can modulate the affinity of the second binding event, thereby influencing ternary complex assembly.^11,12^ This modulation can be either positively or negatively cooperative, depending strongly on the choice of the target protein-binding ligand and the linker length within the PROTAC compound.^13^ The linker length and target protein ligand influence how the POI interacts with the E3 ligase, with residues critical for protein interactions often occupying energetically strained states. PROTACs with strong cooperativity display more such strained contacts at their interfaces than those with weak cooperativity.^14^ In this study, we examined two ligands, JQ1 and I-BET726, which exhibit distinct cooperative behaviors with the target protein bromodomain-containing protein 4 (BRD4).^15^ Specifically, we utilized a truncated construct consisting of the second bromodomain of BRD4 (BRD4^BD2^). JQ1 promotes ternary complex assembly via positive cooperativity, whereas I-BET726 hinders assembly through negative cooperativity.

In addition to BRD4^BD2^ targeting ligands, we also explored the effects of linker length, as prior studies have shown that linker composition can significantly influence cellular activity.^15–17^ When a linker is too short, simultaneous binding of the two proteins becomes challenging due to steric hindrance, thereby impeding ternary complex formation.^18^ If the linker is too long, the distance between the two proteins increases, inhibiting protein-protein interactions and reducing the likelihood that the target protein will be ubiquitinated and marked for degradation.^18^

Native mass spectrometry (nMS) uses electrospray ionization to transfer and kinetically trap biomolecules, including protein and nucleoprotein complexes, into the gas phase, with their non-covalent interactions preserved.^19,20^ This ability makes nMS an attractive technique for studying PROTAC complexes.^21–23^ In nMS, gas-phase dissociation is frequently used to probe structural features of macromolecular complexes, providing insights into subunit connectivity, ligand-binding interactions, and overall topology.^24–26^ Previous research has primarily examined PROTAC-induced ternary systems by isolating the minimal ternary complex to study inter-protein contacts. Recent work has used gas-phase fragmentation to examine PROTAC-mediated complexes, including entire E3 ligase systems, revealing how Cullin-2 stabilizes the VCB core.^27^ In this work, we focus on the ternary complex to examine how changes in the targeting ligand and linker length affect conformation, connectivity, and stability. Surface-induced dissociation (SID) has emerged as a powerful alternative for probing the quaternary structure of protein complexes.^28^ In contrast to other gas-phase fragmentation techniques, SID delivers rapid, high-energy deposition, favoring cleavage of interfaces^28,29^ and yielding subcomplexes whose composition more closely mirrors their native solution-like structure.^30,31^ At lower SID potentials, the weakest non-covalent interfaces are cleaved first.^32^ This selective fragmentation provides valuable insight into the relative strengths of protein-protein interfaces and reveals the connectivity among subunits within a complex.^33^

Here, we employ an integrated nMS approach, combining ion mobility, collision-induced dissociation (CID), SID, and variable-temperature ESI (vT-ESI) to systematically evaluate how linker length and targeting ligand selection govern PROTAC ternary complex topology, interfacial mechanics, and thermal stability. We compared PROTACs sharing the same BRD4^BD2^ targeting ligand but differing in linker length, as well as PROTACs with matched linker lengths but distinct BRD4^BD2^ targeting ligands. Our study focused on seven PROTACs, four containing the JQ1 ligand with linker lengths of 8 (JQ8L), 10 (JQ10L), 13 (JQ13L), and 16 (JQ16L) heavy atoms in the linker, and three containing the I-BET726 ligand with linker lengths of 7 (IBET7L), 10 (IBET10L), and 13 (IBET13L) heavy atoms (Figure S1).

## Results

### Ternary Assembly and Conformational Distributions

The PROTACs used in this study induce ternary complex formation between BRD4^BD2^ and VCB by noncovalently linking V to BRD4^BD2^. As previously shown, BRD4^BD2^ and VCB have not been observed to form a complex without the addition of a PROTAC.^24,27^ Ternary complexes formed by BRD4^BD2^-directed ligands, JQ1 and I-BET726, were investigated using nMS (Figure S2, Figure S3). Under native conditions, the assemblies exhibited narrow, low-charge-state distributions below the theoretical Rayleigh limit, reflecting intact, folded protein topologies.^34^ The measured mass for each of the seven assemblies closely matches the expected mass of the 1:1:1 stoichiometric ternary complex (Figure S2, Table S1). Particularly, signals corresponding to the intact ternary complex dominate the mass spectra across most of the panel, with minimal contributions from unbound protein constituents or binary subcomplexes (Figure S2, Figure S3). This highlights highly efficient assembly under optimized native conditions, except for IBET7L and IBET10L, where ternary assembly is attenuated. For these two variants, while the intact ternary complex is detectable, the mass spectra are instead dominated by the binary species PROTAC-VCB with a minor amount of BRD4^BD2^-PROTAC, alongside free protein constituents (Figure S2E-F, Figure S3E-F). Because all components were mixed under the same conditions, this accumulation of binary intermediates alongside a pronounced decrease in intact ternary complex directly reflects the negative cooperativity and linker length of the I-BET726 constructs, which thermodynamically penalizes secondary binding.

Because gas-phase structure can depend on charge state, we first identified the charge state that best preserves native-like features using surface-induced unfolding (SIU) (Figure S4). Unlike collision-induced unfolding (CIU), SIU drives ions into a large target surface. This high “pressure/temperature jump” energy transfer accesses unique intermediate states that may remain unavailable during the multistep/step-wise activation of CIU, thereby yielding complementary structural insights.^35–37^ Initial screening of the JQ10L-mediated ternary complex across multiple charge states revealed that lower charge states retained compact, native-like structures, whereas higher charge states exhibited elongated profiles prone to premature gas-phase restructuring, consistent with previous work.^38,39^ The 10+ charge state was found to optimally preserve the native-like interfacial contacts of the intact complex under vacuum conditions (higher charge states showed an increase in average CCS and a broadening distribution of CCS values) and 10+ was therefore selected as the baseline for all subsequent structural comparisons (Figure S4).

Having confirmed that all seven PROTAC variants successfully form ternary complex assemblies in solution (Figure S2), we next sought to determine whether these complexes adopt distinct shapes based on their linker lengths. IMS was used to assess the effects of linker length and targeting ligand variation on the topology of the ternary complexes (Figure 2). With the Q-IM-TOF, the 10+, 11+ and 12+ charge states were still observed for each ternary complex (Figure S5). The 10+ charge state was selected for CCS analysis. Each studied complex has at least two distinct conformations, with a prominent conformation at (∼4000 Å^2^) and a minor conformation at (∼3600 Å^2^) (Figure 2). The ∼4000 Å^2^ conformation has previously been observed in a native-like ternary complex.^24^ The predominance of the ∼4000 Å^2^ state suggests that most PROTAC-induced assemblies adopt a conformation consistent with productive E3-target engagement, as productive ternary conformations are directly linked to ubiquitination and degradation efficiency.^40^

**Figure 2:**
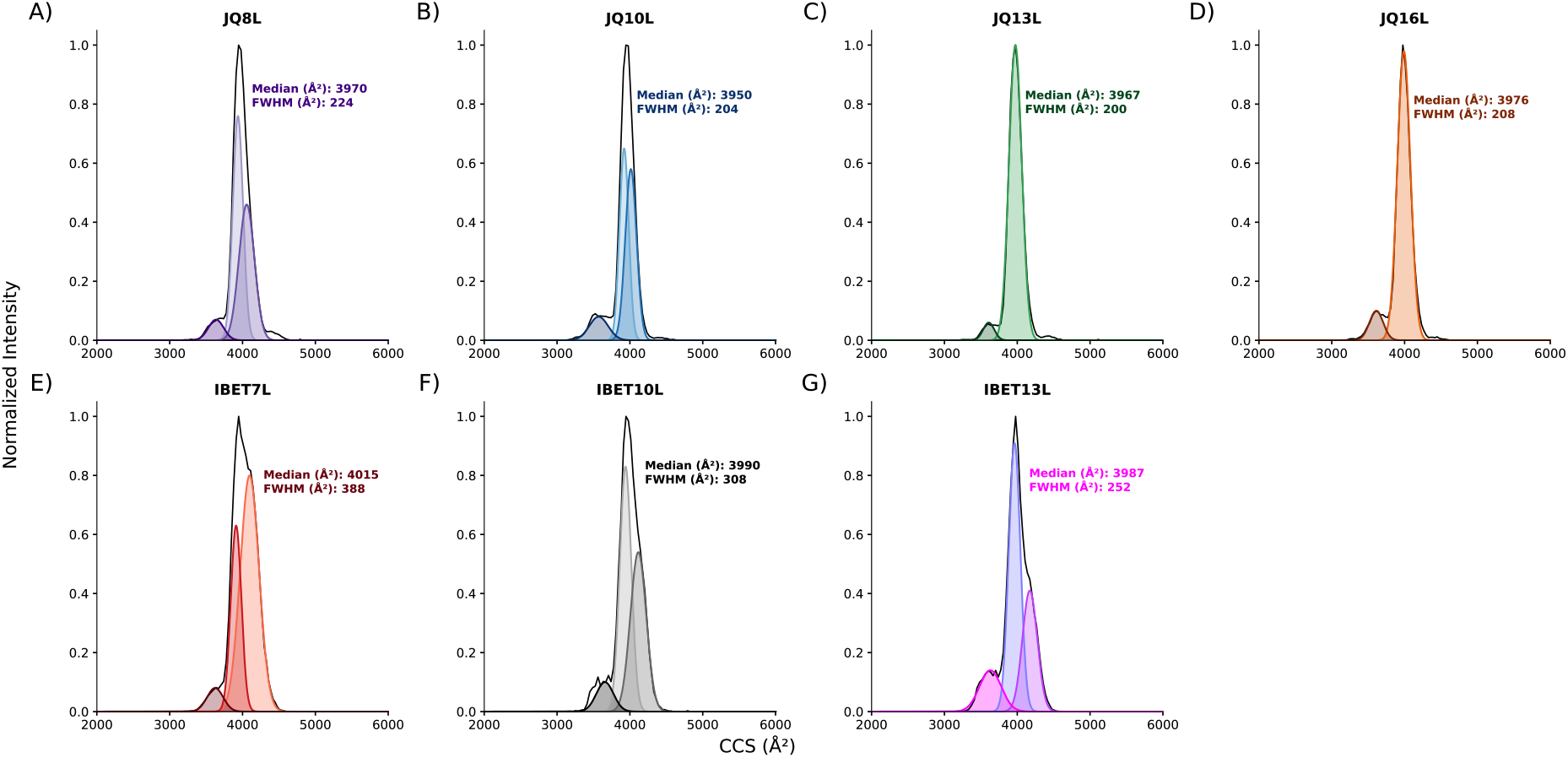
IM-MS analysis of various ternary PROTAC-mediated complexes. CCS distribution of (A). BRD4^BD2^-JQ8L-VCB. (B). BRD4^BD2^-JQ10L-VCB. (C). BRD4^BD2^-JQ13L-VCB. (D). BRD4^BD2^-JQ16L-VCB. (E). BRD4^BD2^-IBET7L-VCB. (F). BRD4^BD2^-IBET10L-VCB. (G). BRD4^BD2^-IBET13L-VCB.

Longer linkers allow the ternary complexes to adopt energetically favorable conformations more easily.^41^ This is observed through a stable, consistent ternary topology. Although CCS values increase with mass, the narrower, more uniform distribution of longer linker lengths (JQ13L, JQ16L, IBET10L, IBET13L) suggests a more narrowly distributed, reproducible conformation (Figure 2, Table S2). This flexibility in the PROTAC molecule may facilitate dynamic sampling of conformations until it reaches the most favorable geometry.

By contrast, for shorter linker lengths, the centroids of the CCS distributions (3970 Å^2^ and 4015 Å^2^) indicate that rigid linkers force the ternary complex into more extended shapes. This may reflect a structural difference in engagement, in which reduced PROTAC flexibility in shorter-linker complexes can impose conformational restraints. Such conformations may reduce productive interface formation between BRD4^BD2^ and the E3 ligase, lowering degradation efficiency.^15^ Overly rigid linkers may hinder the conformational flexibility required during ternary complex formation, potentially reducing the efficiency or likelihood of complex assembly.^42^ Steric strain may drive the complex toward more extended or rigid topologies, thereby increasing its overall size and surface area and resulting in an elevated CCS.

The JQ1 series forms a predominantly single, compact conformation centered at ∼4000 Å^2^ with narrow distribution widths (FWHM ∼200-224 Å^2^), whereas the I-BET726 series exhibits greater conformational heterogeneity, populating two distinct states (∼3900 Å^2^ and ∼4100 Å^2^) with broader distributions (FWHM ∼252-388 Å^2^) (Figure 2). Previous work has shown the JQ1 series of PROTACs forms a highly dynamic, flexible ternary complex that can sample many different physical conformations, whereas the I-BET726 series is much more structurally restricted, locking the ternary complex into fewer possible shapes.^43^ The inherent flexibility of JQ1 complexes allows them to collapse into compact conformations, whereas rigid I-BET726 complexes are trapped across multiple non-relaxed structural states.

Mechanistically, this heterogeneity could reflect alternative binding orientations of BRD4^BD2^ relative to V. Such heterogeneity could affect degradation efficiency, as only conformations that optimally position the substrate for ubiquitin transfer are likely to be functional. It has been demonstrated that the JQ1 targeting ligand series is a more effective overall degrader than I-BET726,^15^ which may indicate a more uniform structural ensemble. These findings suggest that the scaffold’s steric environment can influence the conformational landscape of the ternary complex, with I-BET726 introducing greater heterogeneity, whereas JQ1 favors a dominant, compact arrangement.

### Comparative Stability via Energy-Resolved CID

CID gradually deposits internal energy through repeated collisions with neutral gas molecules, often resulting in structural rearrangement and unfolding before dissociation along low-energy pathways.^28^ Energy-resolved CID can therefore provide comparative measures of gas-phase complex stability and interaction strength in cases where dissociation outcompetes restructuring.^44,45^ For previous PROTAC ternary complexes studied, the primary dissociation pathway by CID is PROTAC ejection.^24,26,27^

To compare the energy required for PROTAC release, the 10+ charge state of each ternary complex was isolated and subjected to a range of CID energies (Figure S6). Relative gas-phase stability was assessed using CID_50_, defined as the collision energy required to reduce the precursor abundance to 50% of its initial value (Figure 3).

**Figure 3:**
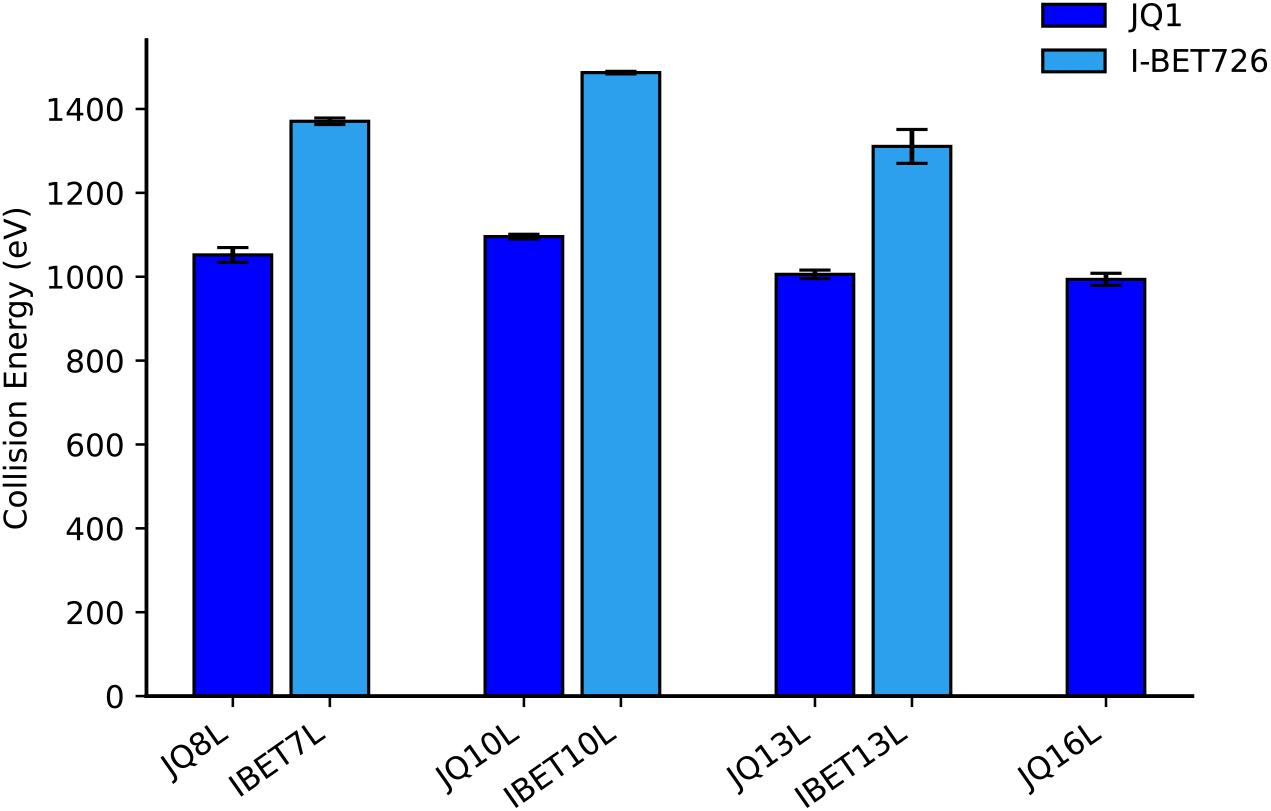
CID_50_ analysis of each PROTAC-mediated complex studied. CID_50_ is defined as the CID collision energy required to reduce the precursor to 50% of its original amount.

CID_50_ values showed no systematic dependence on linker length in either the JQ1 or I-BET726 series, indicating that linker length does not determine the energy required for PROTAC ejection. Instead, dissociation energetics depended primarily on the targeting ligand: I-BET726-based complexes consistently required higher collision energies for PROTAC release than JQ1-based complexes. Because all variants contained the same protein partners, the higher CID_50_ values of the I-BET726 complexes likely reflect stronger local binding within the BRD4^BD2^ pocket.

### Dissecting Interfacial Connectivity via SID

Unlike the slow, multistep activation used in CID, SID deposits energy in a single high-energy collision, limiting unfolding before dissociation and providing a more direct probe of noncovalent interfaces. All seven ternary complexes showed broadly similar classes of SID products (Figure 4 and Figure S7). However, SID_50_ values were generally higher for the I-BET726-based complexes than for the JQ1-based complexes (Figure S8), whereas the 10+ precursor ions showed similar surface-induced unfolding behavior (Figure S9). These results indicate that the complexes access similar fragmentation pathways but differ in the energy required for dissociation.

**Figure 4:**
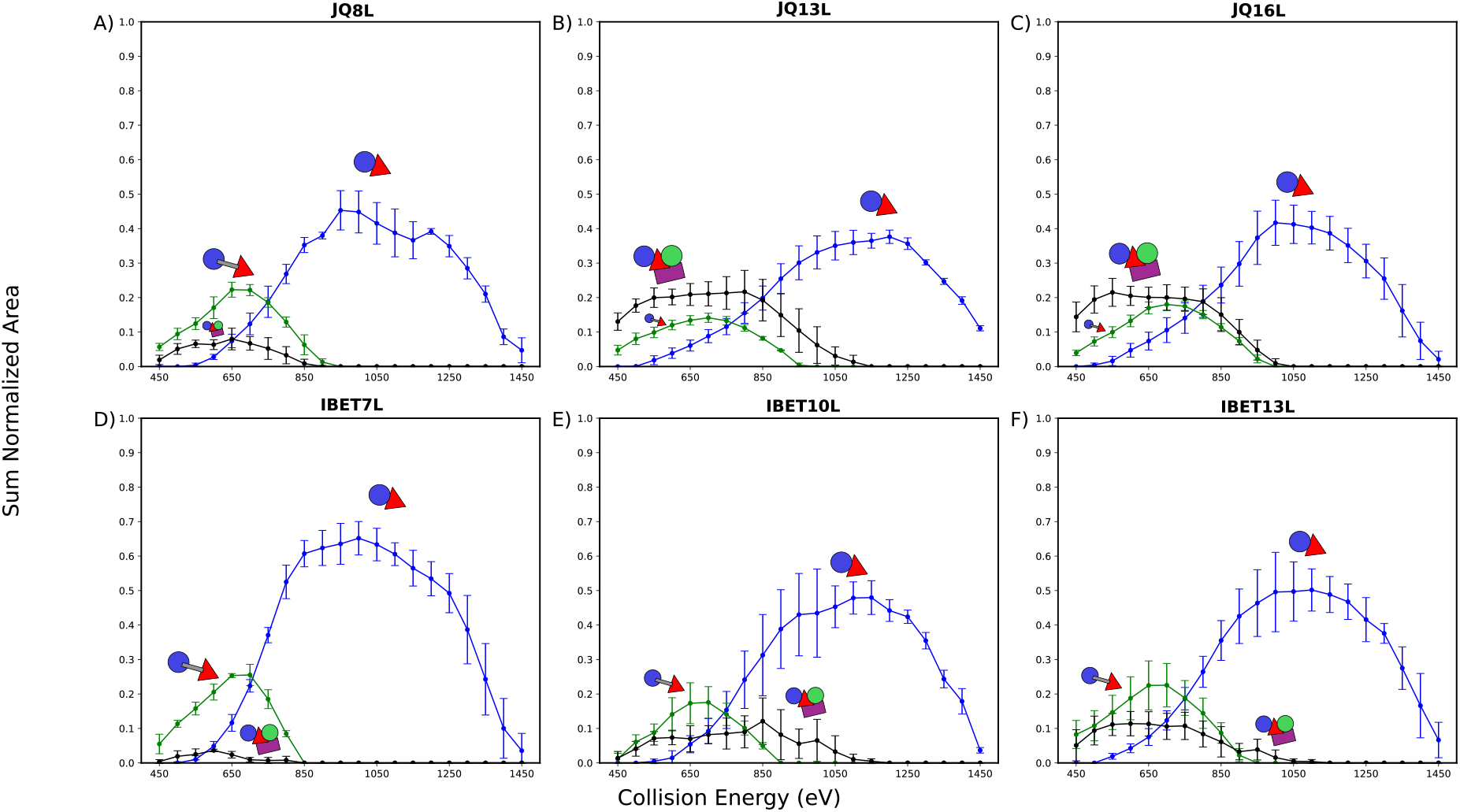
ERMS plots for 10+ BRD4^BD2^-PROTAC-VCB. The fragments shown correspond to BRD4^BD2^-VCB (PROTAC ejection), BRD4^BD2^-PROTAC-V (CB loss), and BRD4^BD2^-V. For clarity, selected fragments have been omitted. See SI Figure S7 for further data on all dissociation products. (A). JQ8L. (B). JQ13L. (C). JQ16L. (D). IBET7L. (E). IBET10L. (F). IBET13L. (JQ10L behavior was reported previously.^27^)

Within the JQ1 series, JQ8L produced a distinct product distribution. In contrast to the longer-linker JQ1 complexes, JQ8L showed minimal PROTAC ejection and a higher relative abundance of BRD4^BD2^-PROTAC-V than BRD4^BD2^-VCB following SID (Figure 4A). Comparisons across targeting ligands revealed systematic differences in PROTAC retention. With the exception of JQ8L, the JQ1-based complexes predominantly formed BRD4^BD2^-VCB through PROTAC ejection, consistent with previous results for JQ10L.^27^ In contrast, the I-BET726-based complexes produced more BRD4^BD2^-PROTAC-V than BRD4^BD2^-VCB at low SID energies (450-650 eV; Figure 4D-F), indicating greater retention of the BRD4^BD2^-PROTAC interface. This trend is consistent with the higher CID_50_ values measured for the I-BET726 complexes (1310-1480 eV) than for the JQ1 complexes (∼1000 eV; Figure 3). Together, the SID and CID data show that targeting-ligand identity has a stronger effect than linker length on PROTAC retention during gas-phase activation, with JQ8L representing a distinct case within the JQ1 series.

### Conformational Resiliency of PROTAC Ejected Subcomplexes

To assess whether JQ10L ejection induces substantial conformational changes, we measured the collision cross sections (CCS) of SID-generated subcomplexes and compared them with those of the corresponding JQ10L-bound species (Figure 5). The 9+ BRD4^BD2^-VCB subcomplex exhibited a compact conformation centered at ∼3700 Å^2^, compared with ∼3800 Å^2^ for the compact 10+ BRD4^BD2^-JQ10L-VCB precursor, corresponding to a 1.8% decrease in mass and a 2.7% decrease in CCS (Figure 5A). Similarly, BRD4^BD2^-JQ10L-V and BRD4^BD2^-V exhibited CCS values of ∼2700 and ∼2650 Å^2^, respectively; JQ10L ejection produced a 2.9% decrease in mass and a ∼2.0% decrease in CCS (Figure 5B). JQ10L-V and V showed CCS values of ∼1900 and ∼1850 Å^2^, respectively, corresponding to a 5.2% decrease in mass and a 2.6% decrease in CCS (Figure 5C). Across all three comparisons, JQ10L ejection produced only small reductions in CCS, consistent with modest compaction of the remaining protein subcomplexes rather than extensive structural rearrangement.

**Figure 5:**
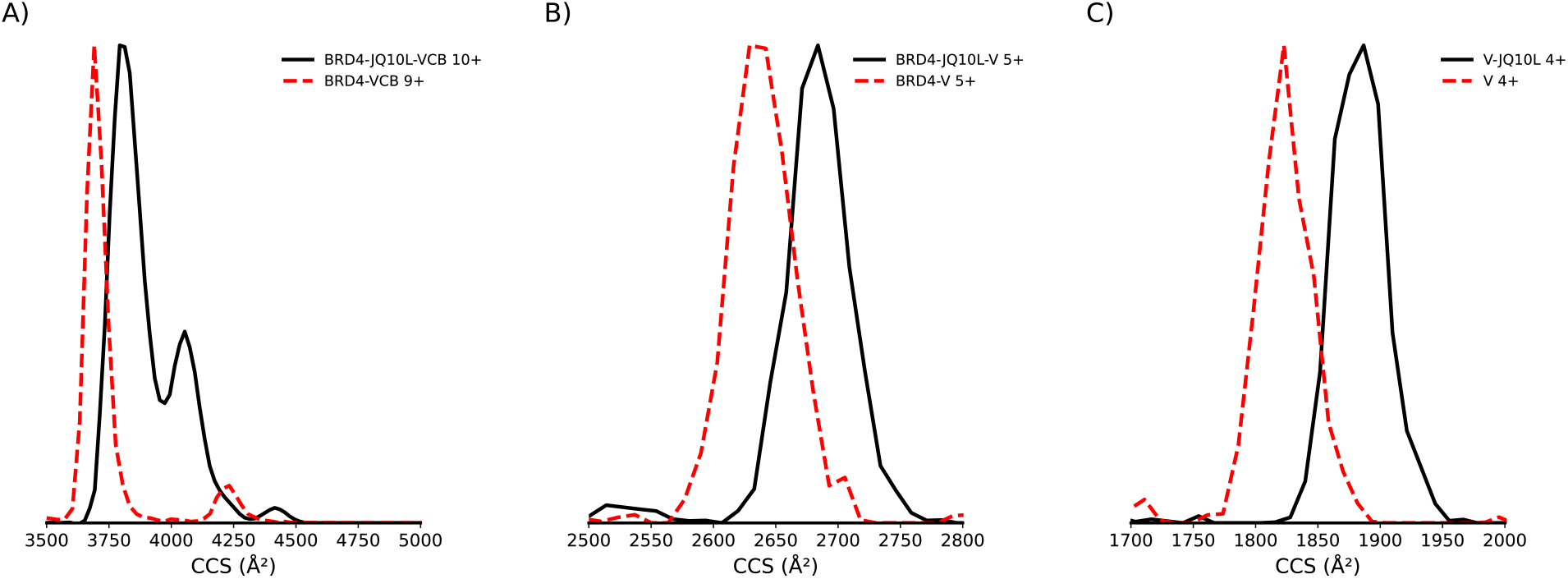
CCS distributions of SID-generated products of 10+ BRD4-JQ10L-VCB at SID 80 V. (A). CCS distributions of BRD4^BD2^-JQ10L-VCB and BRD4^BD2^-VCB. (B). CCS distributions of BRD4^BD2^-JQ10L-V and BRD4^BD2^-V. (C). CCS distributions of JQ10L-V and V.

Overall, SID provides detailed insight into the conformational landscape of the JQ10L-mediated complex and its subcomponents. Minimal CCS reductions after JQ10L ejection suggest that the protein scaffolds remain intact, while JQ10L ejection leads to slightly more compact structures, which is likely to be primarily due to the loss of mass. The overall architecture remains largely preserved, supporting the idea that JQ10L ejection does not substantially alter the complex’s core structural framework.

### Comparative Stability via vT-ESI

To bridge our kinetically controlled gas-phase fragmentation insights with solution-phase thermodynamics, we evaluated the thermal stabilities and equilibrium behaviors of the ternary complexes using variable-temperature ESI (vT-ESI). These measurements focused on JQ10L- and IBET10L-mediated complexes, which share the same linker length but have distinct BRD4^BD2^ targeting ligands. The JQ1-based PROTAC, JQ10L, displayed a single monophasic melting transition with a Tm of ∼50 °C (Figure 6A). In contrast, the I-BET726-based PROTAC, IBET10L, exhibited a distinct biphasic melting curve with transitions at ∼35 °C and ∼45 °C (Figure 6B). This biphasic melting behavior for IBET10L may be explained by the conformational heterogeneity (and dissociation of one of the populations; Figures S10-12) observed in our ion mobility CCS distributions, in which IBET10L populates both compact 3900 Å^2^ and extended 4100 Å^2^ states (Figure 2F).

**Figure 6:**
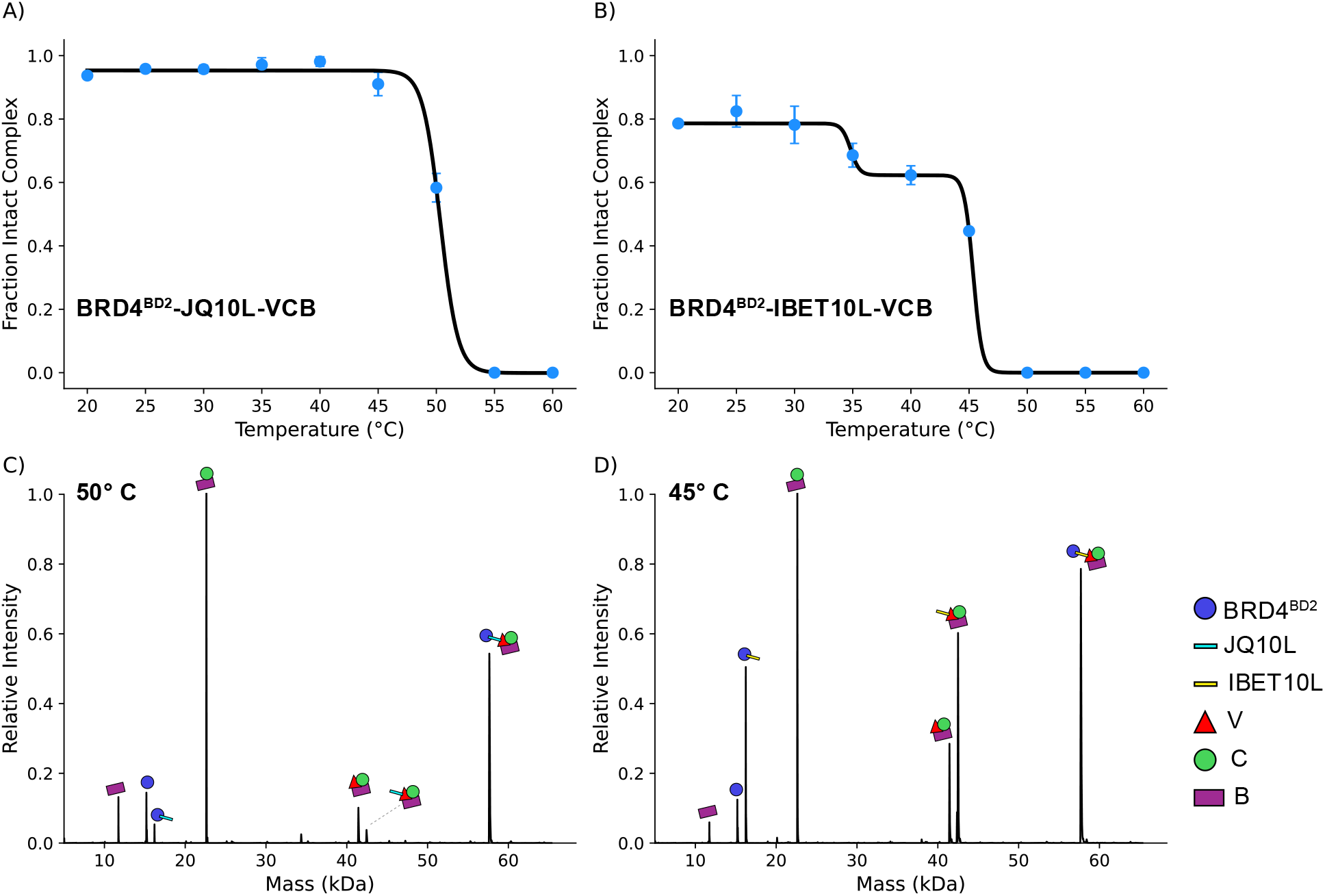
vT-ESI analysis for (A) BRD4^BD2^-JQ10L-VCB and (B) BRD4^BD2^-IBET10L-VCB ternary complexes. Curves depict nonlinear regression using a sigmoidal Boltzmann equation. (C). Mass distribution of BRD4^BD2^-JQ10L-VCB sprayed at 50 °C with n=3 replicates. (D). Mass distribution of BRD4^BD2^-IBET10L-VCB sprayed at 45 °C with n=2 replicates. In (B), the lower initial fraction is the result of a higher proportion of free components present (see Figures S2, S3, and S12).

## Discussion and Conclusions

This study uses complementary native MS measurements to determine how linker length and BRD4-targeting-ligand identity influence PROTAC-induced ternary complexes. The results distinguish effects on conformational distributions, gas-phase dissociation behavior, interfacial retention, and solution-phase thermal disassembly.

I-BET726-based complexes populated prominent states near 3900 Å^2^ (minor) and 4100 Å^2^, while JQ1-based complexes predominantly populated a state near 4000 Å^2^. Thus, linker length and targeting-ligand scaffold influenced the observed conformational distributions, although these data do not establish whether individual conformations are productive for ubiquitination or degradation.

CID_50_ values did not vary systematically with linker length, but I-BET726-based complexes required higher collision energies for PROTAC ejection than JQ1-based complexes. In SID, I-BET726 complexes preferentially retained BRD4^BD2^-PROTAC-V at low activation energies, whereas most JQ1 complexes predominantly formed BRD4^BD2^-VCB through PROTAC ejection. JQ8L was an exception within the JQ1 series, producing little PROTAC ejection and a greater relative abundance of BRD4^BD2^-PROTAC-V. These results indicate that targeting-ligand identity had a stronger effect than linker length on PROTAC retention during gas-phase activation.

CCS measurements of SID-generated JQ10L subcomplexes showed decreases of approximately 2.7% for BRD4^BD2^-VCB, 2.0% for BRD4^BD2^-V, and 3.4% for V relative to their JQ10L-containing counterparts. These modest reductions in CCS associated with PROTAC loss suggest that JQ10L ejection did not substantially disrupt the architecture of the detected protein subcomplexes.

vT-ESI measurements revealed distinct thermal behavior for the matched-linker JQ10L and IBET10L complexes. JQ10L showed a single transition near 50 °C and released V and CB, whereas IBET10L exhibited transitions near 35 and 45 °C and accumulated VCB and IBET10L-VCB before V increased at the second transition. The biphasic behavior of IBET10L is consistent with multiple structural populations, although the measurements do not establish a direct correspondence between individual CCS populations and thermal transitions.

The different products observed by vT-ESI and SID reflect their distinct activation regimes. vT-ESI permits solution-phase thermal equilibration and structural rearrangement before dissociation, whereas SID rapidly activates isolated gas-phase ions and reports on interfaces retained during dissociation. To demonstrate, vT-ESI was applied to streptavidin (a protein with a known dimer-of-dimers topology), generating only monomers from the tetramer upon thermal dissociation in solution (Figure S13). The higher thermal transition of JQ10L is consistent with its reported greater cellular degradation efficacy relative to IBET10L,^15^ as a more stable ternary assembly can facilitate more productive target ubiquitination, but comparison of two matched-linker complexes is insufficient to establish thermal behavior as a general predictor of degradation efficacy.

Overall, linker length primarily influenced the conformational distributions observed by ion mobility, whereas targeting-ligand identity more strongly affected PROTAC retention during CID and SID. The differing behaviors observed during gas-phase activation and solution-phase thermal disassembly demonstrate that no single metric fully captures ternary-complex behavior. Integrating these measurements therefore provides a clearer basis for determining how PROTAC architecture influences complex conformation, interfacial connectivity, and disassembly.

## Materials and Methods

### Sample Preparation

PROTACs, BRD4^BD2^, and the VCB complex were provided by AbbVie. Proteins were expressed and purified as previously described.^27^ Proteins were buffer-exchanged into 200 mM ammonium acetate (AmAc) using Bio-Rad 6 kDa size-exclusion spin columns. PROTAC stock solutions were prepared in DMSO and diluted with water to 100 µM PROTAC and 1% (v/v) DMSO before addition to the protein samples.^21^ After buffer exchange, BRD4^BD2^, PROTAC, and VCB were combined at a 1:2:1 molar ratio (5 µM BRD4^BD2^, 10 µM PROTAC, and 5 µM VCB) and incubated at room temperature for 15 min to form the ternary complex. For SID experiments, triethylammonium acetate (TEAA) was added to a final concentration of 20% (v/v) to reduce the charge states of the complexes.^30^

### Mass Spectrometry

#### nMS and SID-ERMS

Native MS, CID, and SID experiments were performed on a Q Exactive™ UHMR Hybrid Quadrupole Orbitrap™ mass spectrometer (Thermo Scientific, Bremen, Germany) modified with a 4-cm SID device.^46^ Samples (3-5 µL) were loaded into in-house-pulled borosilicate capillaries prepared with a P-97 micropipette puller (Sutter Instruments, Novato, CA) and ionized by nanoelectrospray ionization in positive-ion mode. Typical source and acquisition settings were as follows: spray voltage, 0.5-0.9 kV; capillary temperature, 250 °C; in-source trapping, −30 V; trap gas flow setting, 5, corresponding to an ultrahigh-vacuum gauge reading of approximately 4.3 × 10^−10^ mbar; and resolving power, 12,000. For SID-ERMS measurements, the 10+ ternary-complex precursor was isolated with the quadrupole and subjected to SID potentials of 45-145 V; additional instrument settings are provided in Table S3. For CID-ERMS measurements, the 10+ precursor was isolated and activated over an experiment-specific collision-energy range [50-1950 eV with increments of 50 eV]. Precursor and product abundances were normalized at each activation energy, and CID_50_ or SID_50_ was defined as the energy at which precursor abundance decreased to 50% of its initial value [Collision energy values corresponding to 50% normalized intensity]. [CID_50_/SID_50_ values were determined via 1D linear inverse interpolation of the experimental data points using the interp1d function (scipy.interpolate) in Python with 3 replicates]. Thermal-dissociation experiments were performed using an in-house-built variable-temperature electrospray ionization (vT-ESI) source interfaced with the same mass spectrometer, as described previously.^47^ Samples were equilibrated for 2 min at each temperature [20-60 °C with 5 °C intervals]. Relative abundances of the ternary complex and dissociation products were determined from deconvolved spectra using UniDec.^48^ [ *m/z* range: 1000-10000; charge range: 1-20; mass range: 5000-65000; peak FWHM (Th): 3; peak shape function: Gaussian; Beta: 0; charge smooth width: 1.0; point smooth width: 1.0; mass smooth width: 0.0. Melting curves were fit by weighted non-linear least-squares regression (scipy.optimize.curve_fit). JQ10L datasets (n=3) were fitted to a four-parameter Boltzmann sigmoidal model. IBET10L datasets (n=2) were fitted to a biphasic double-sigmoidal model, where transition temperatures were identified from local maxima in the negative first derivative using SciPy’s peak-detection algorithm]. Mass spectra were extracted and analyzed using Xcalibur 4.1 (Thermo Scientific).

#### nMS and Ion Mobility / CCS Measurements

Ion-mobility experiments were performed on a Waters SELECT SERIES Cyclic IMS (Q-IM-TOF) equipped with a commercially available stainless-steel SID device (3 mm along the ion path). The device was positioned after the quadrupole, in place of the DRE lens and before the trap stacked-ring ion guide used for CID.^35^ Analyses were performed in positive-ion mode using the following settings: capillary voltage, 0.5-0.9 kV; cone voltage, 20 V; source temperature, 25 °C; and N_2_ trap gas flow rate, 10.0 mL/min. Additional cyclic ion-mobility settings are provided in Table S4. Mass spectra were analyzed using MassLynx and DriftScope, and ion-mobility profiles were processed using CIUSuite 3.^49^

Collision cross sections (CCSs) were calibrated using IMSCal^50^ and a reference CCS database^51^ according to established procedures for ion mobility experiments. The calibrant mixture contained β-lactoglobulin monomer, carbonic anhydrase, β-lactoglobulin dimer, streptavidin, and C-reactive protein. Calibrants were buffer-exchanged as described above and analyzed in 160 mM ammonium acetate containing 40 mM TEAA.

## Author Contributions

E.N.W. conducted and examined nMS experiments. J.D.G. and M.M.K. initiated and designed experiments. M.A.E. synthesized the I-BET726-based PROTACs. A.C.E. and T.A.T. expressed and purified BRD4 and VCB. V.H.W. and S.M.M. designed and led the project. All authors contributed to writing the manuscript.

## Data Availability Statement

Supporting data for the findings reported in this manuscript are provided in the Supplementary Material.

## Acknowledgments

This work was supported by National Institutes of Health award RM1GM149374 (to V.H.W.). E.N.W. received funding from an NIH T32 grant at Ohio State (GM144293), and a graduate fellowship from the Georgia Tech School of Chemistry and Biochemistry backed by a U.S. Department of Education GAANN award (P200A240072). Microsoft CoPilot was used to refine grammar and clarity.

## Supporting Information

Additional supporting information can be found online in the Supporting Information section at the end of this article.

