## Supplementary Material for "Native Mass Spectrometry Reveals Distinct Effects of Linker Length and Targeting-Ligand Identity on PROTAC Ternary Complexes"

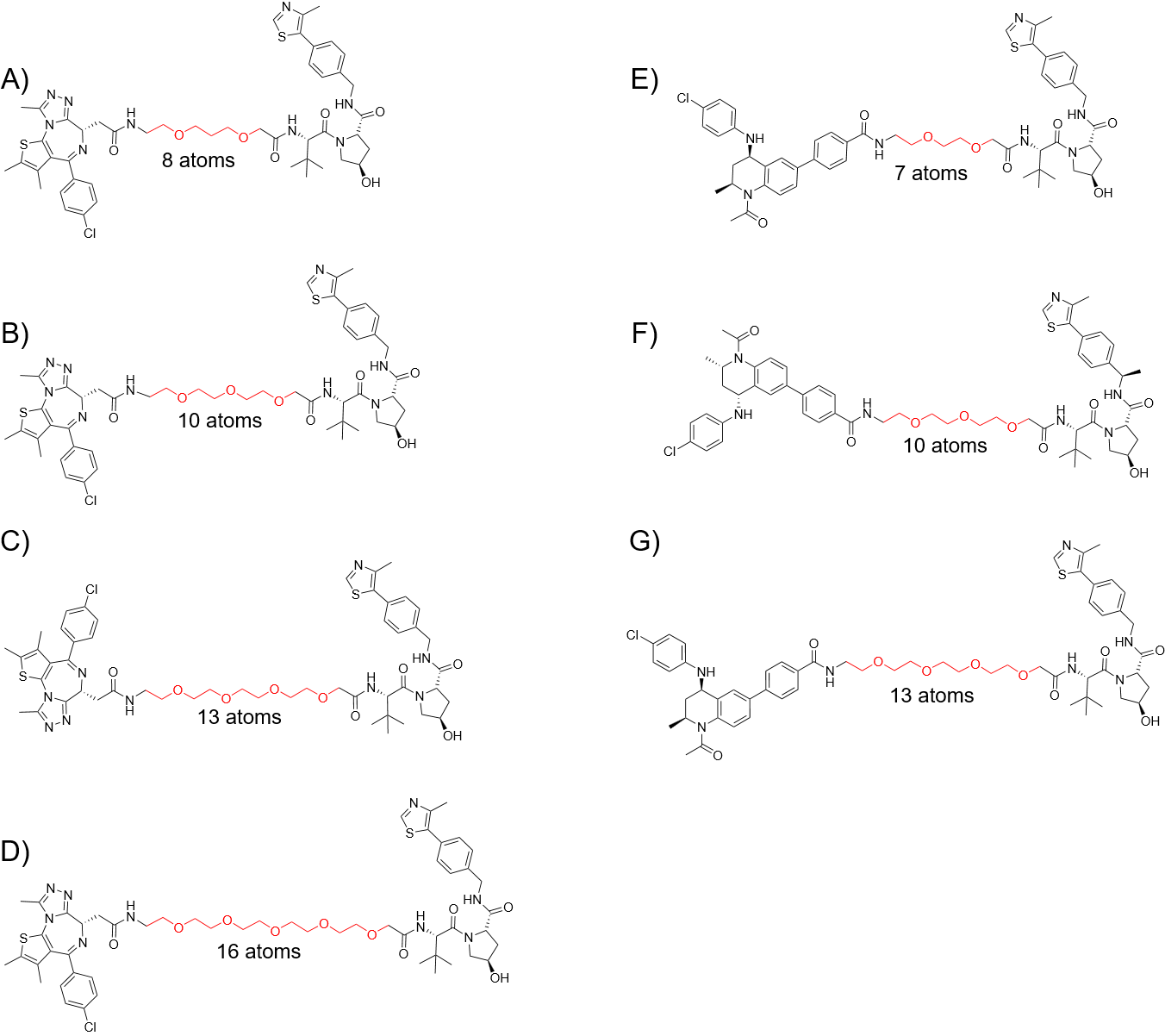

**Figure S1.** Structures of various PROTACs used. A) JQ8L. B) JQ10L (MZ1). C) JQ13L (MZ2). D) JQ16L. E) IBET7L (MZP-61). F) IBET10L (MZP-54). G) IBET13L (MZP-55). Linker lengths correspond to number of heavy atoms in red.

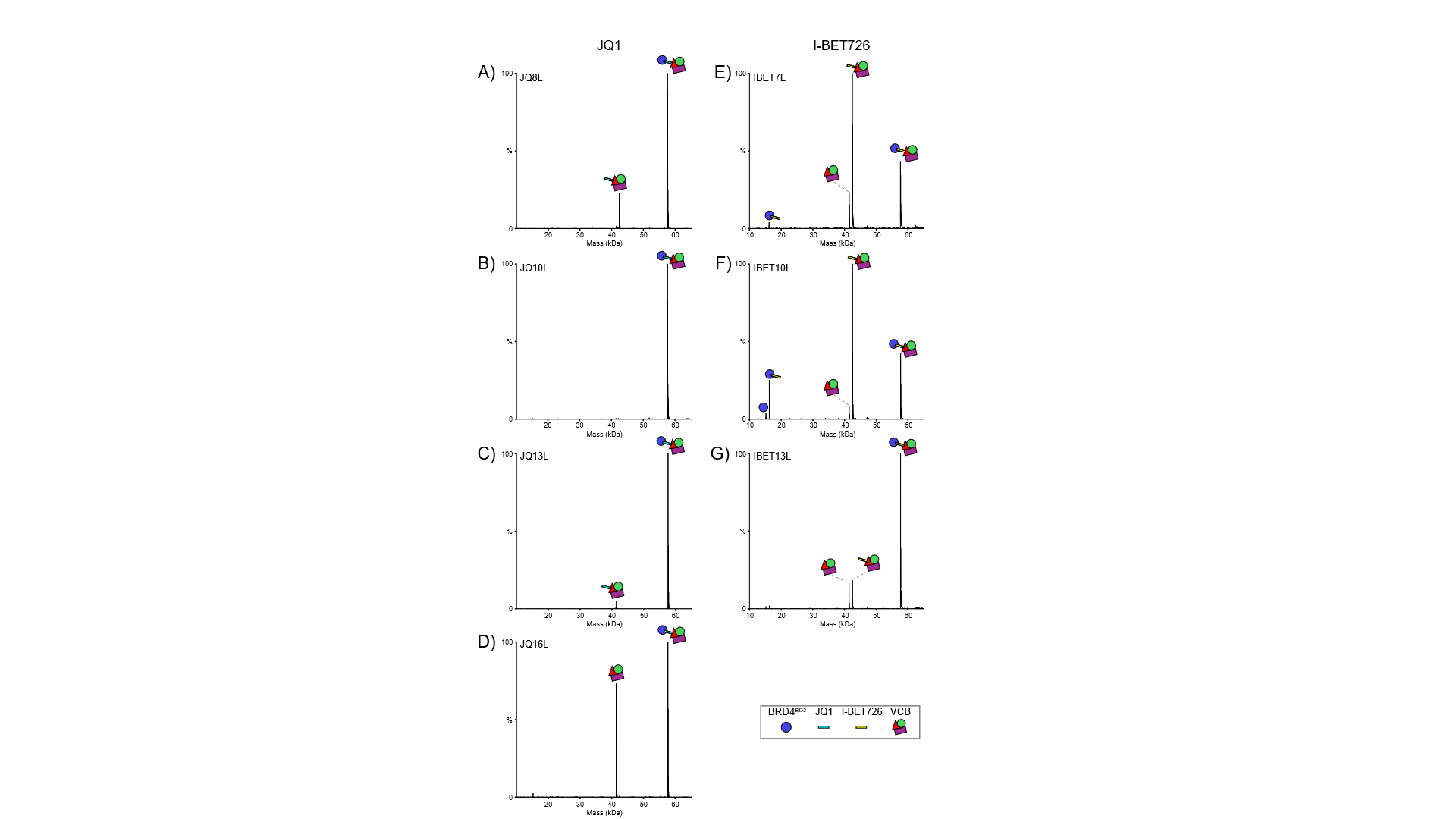

**Figure S2:** nMS mass distributions of 1:2:1 BRD4^BD2^ PROTAC-VCB ternary complexes. Distributions are shown for complexes containing (A-D) JQ1 series PROTACs (JQ8L, JQ10L, JQ13L, JQ16L) and (E-G) IBET-726 series PROTACs (IBET7L, IBET10L, IBET13L).

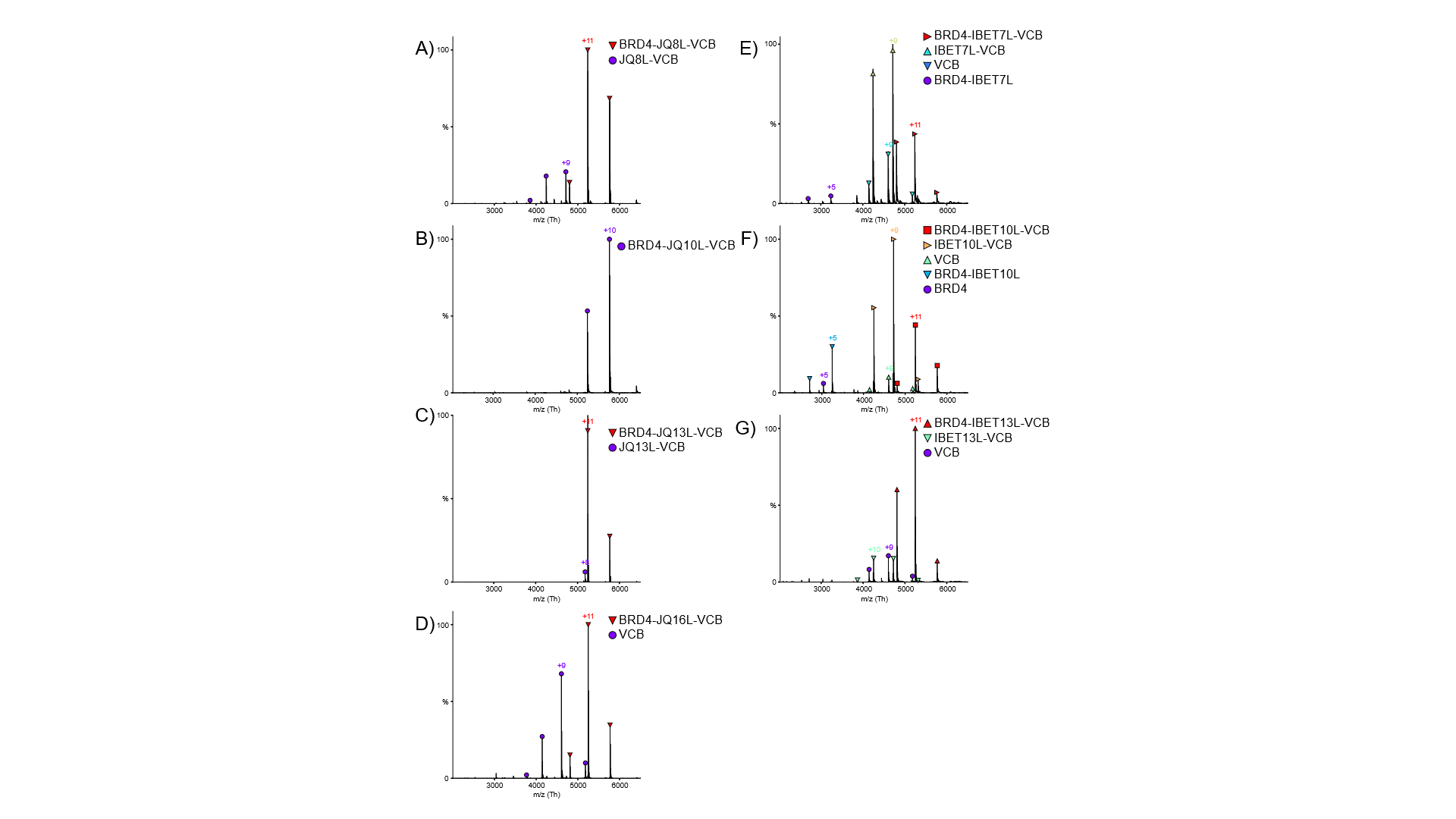

**Figure S3.** nMS spectra of 1:2:1 BRD4^BD2^ PROTAC-VCB ternary complexes. Spectra are shown for complexes containing (A-D) JQ1 series PROTACs (JQ8L, JQ10L, JQ13L, JQ16L) and (E-G) IBET-726 series PROTACs (IBET7L, IBET10L, IBET13L).

| Ternary complex | Measured Mass (Da) | Theoretical Mass (Da) |
| --- | --- | --- |
| JQ8L | 57,599 ±1 | 57,600 |
| JQ10L | 57,629 ±1 | 57,630 |
| JQ13L | 57,674 ±1 | 57,674 |
| JQ16L | 57,718 ±1 | 57,718 |
| IBET7L | 57,622 ±1 | 57,620 |
| IBET10L | 57,680 ±1 | 57,679 |
| IBET13L | 57,711 ±1 | 57,708 |

**Table S1:** Measured and theoretical masses for each ternary complex studied

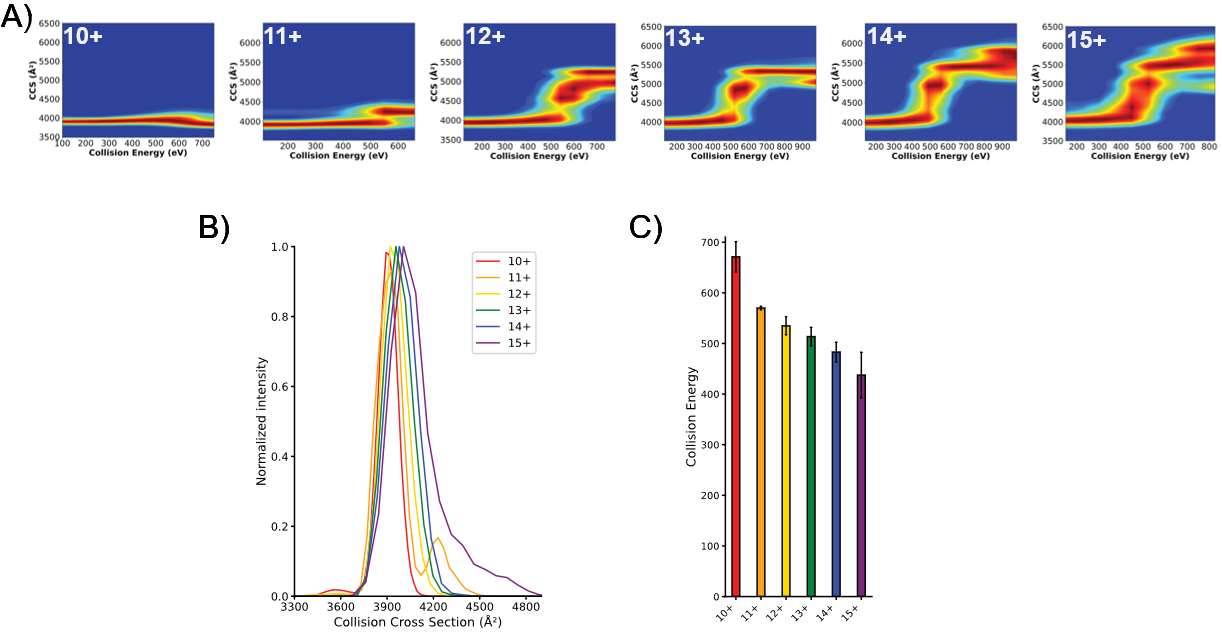

**Figure S4:** IM-nMS analysis of BRD4^BD2^-JQ10L-VCB. (A). Average (n=3) SIU fingerprint for 10-15+ BRD4^BD2^-JQ10L-VCB. (B). CCS distributions of 10-15+ BRD4^BD2^-JQ10L-VCB. (C). SIU_50_ analysis of 10-15+ BRD4^BD2^-JQ10L-VCB.

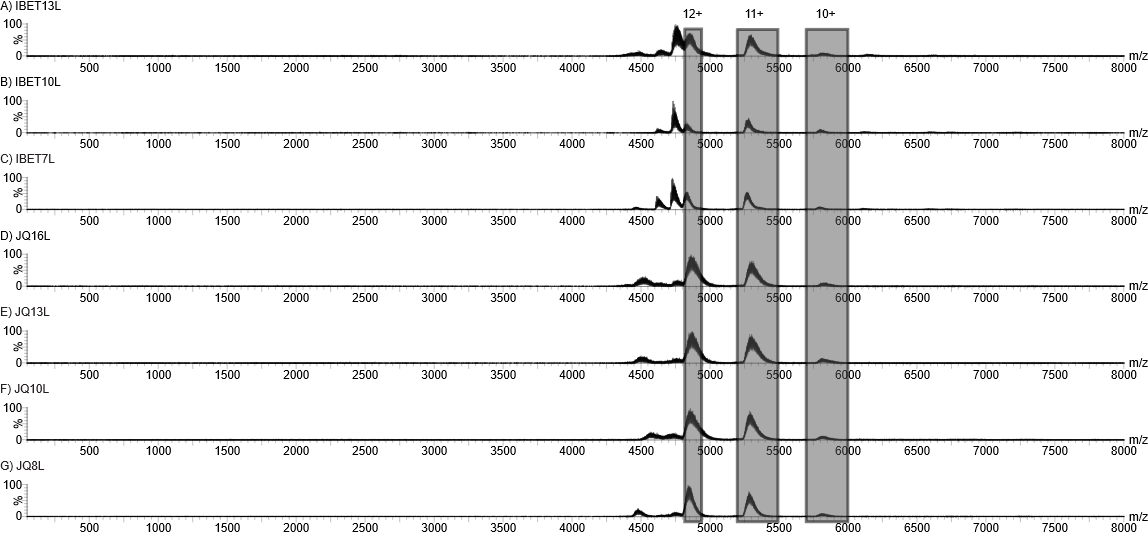

**Figure S5:** nMS analysis of various ternary PROTAC-mediated complexes. (A). MS1 of 1:2:1 BRD4^BD2^-IBET13L-VCB. (B). MS1 of 1:2:1 BRD4^BD2^-IBET10L-VCB. (C). MS1 of 1:2:1 BRD4^BD2^-IBET7L-VCB. (D). MS1 of 1:2:1 BRD4^BD2^-JQ16L-VCB. (E). MS1 of 1:2:1 BRD4^BD2^-JQ13L-VCB. (F). MS1 of 1:2:1 BRD4^BD2^-JQ10L-VCB. (G). MS1 of 1:2:1 BRD4^BD2^-JQ8L-VCB.

| PROTAC | Apex (Å^2^) | Median (Å^2^) | FWHM (Å^2^) |
| --- | --- | --- | --- |
| JQ8L | 3954 | 3970 | 224 |
| JQ10L | 3951 | 3950 | 204 |
| JQ13L | 3974 | 3967 | 200 |
| JQ16L | 3990 | 3976 | 208 |
| IBET7L | 3945 | 4015 | 388 |
| IBET10L | 3959 | 3990 | 308 |
| IBET13L | 3970 | 3987 | 252 |

**Table S2:** Apex, Median and FWHM of summed fit Gaussian distributions of 10+ ternary complexes.

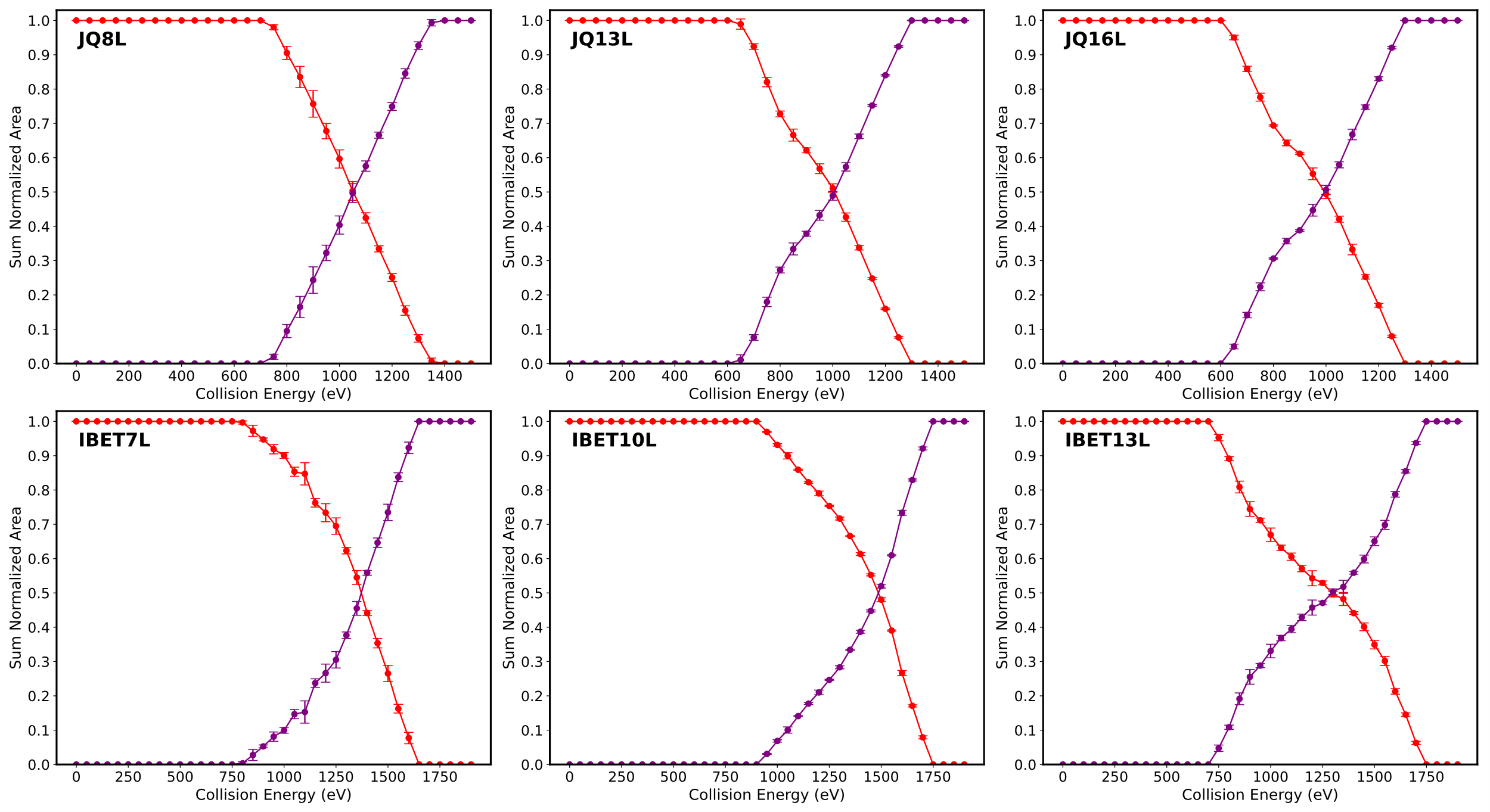

**Figure S6.** CID-ERMS plots for the 10+ charge state of each PROTAC complex studied. The red line depicts the BRD4-PROTAC-VCB precursor and purple line depicts BRD4-VCB fragment which is observed after ejection of the PROTAC. Larger amounts of energy (~300 eV) are required to reach the intersection between the two species for the I-BET726 warhead (bottom row) than the JQ1 warhead (top row). (JQ10L behavior was reported previously.^27^)

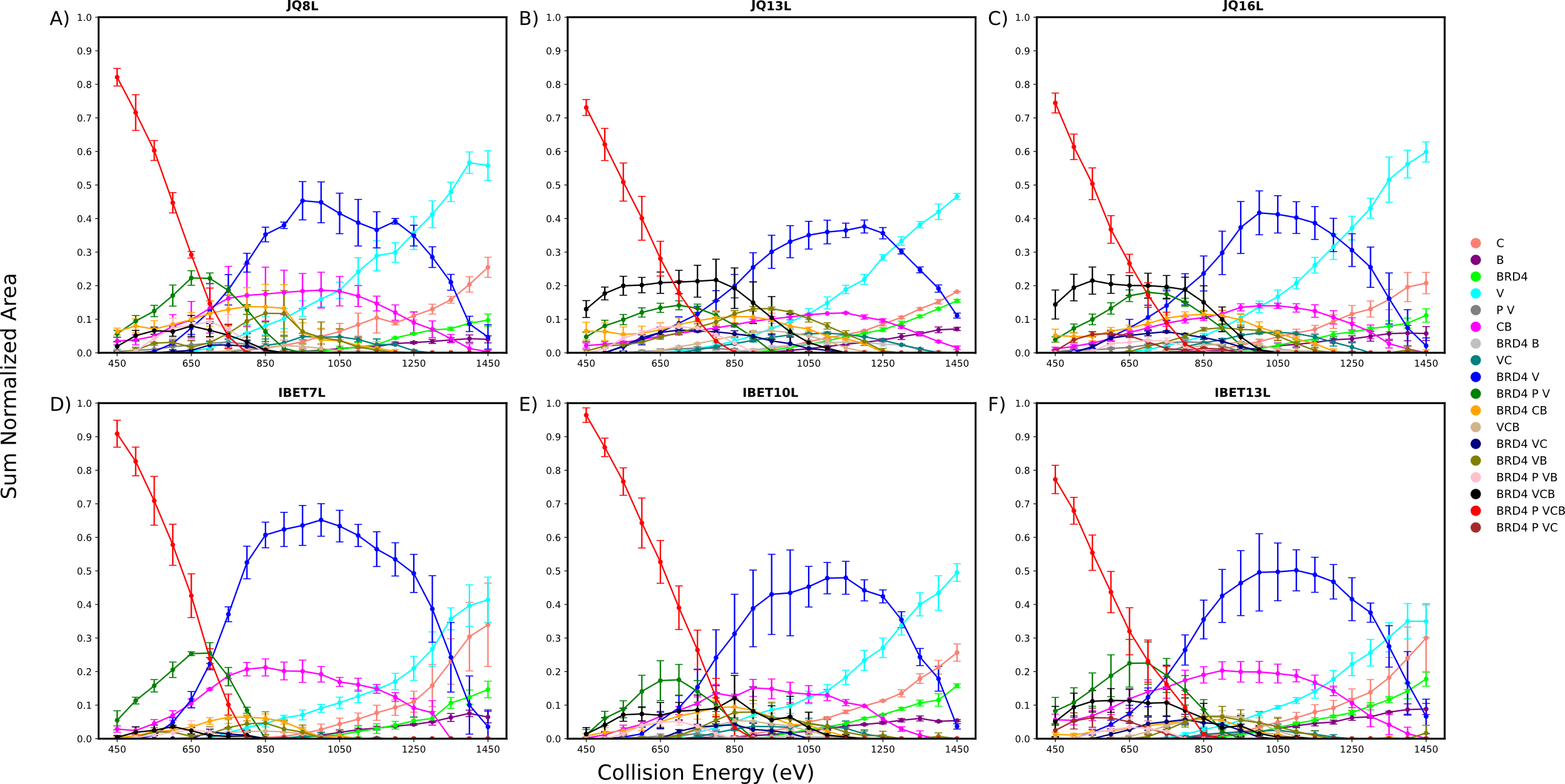

**Figure S7.** SID-ERMS plots for the 10+ charge state of each PROTAC complex studied.

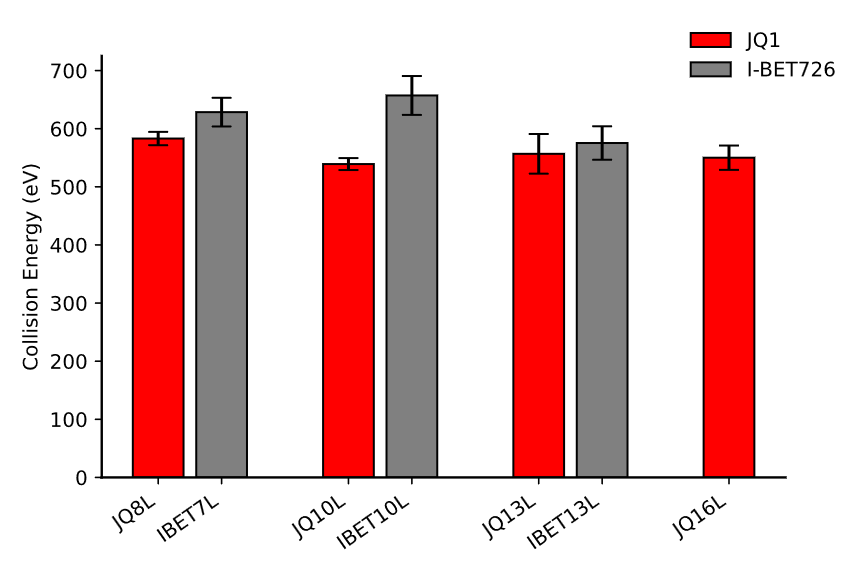

**Figure S8.** SID_50_ analysis of each PROTAC-mediated complex studied. SID_50_ is defined as the SID collision energy required to reduce the precursor to 50% remaining.

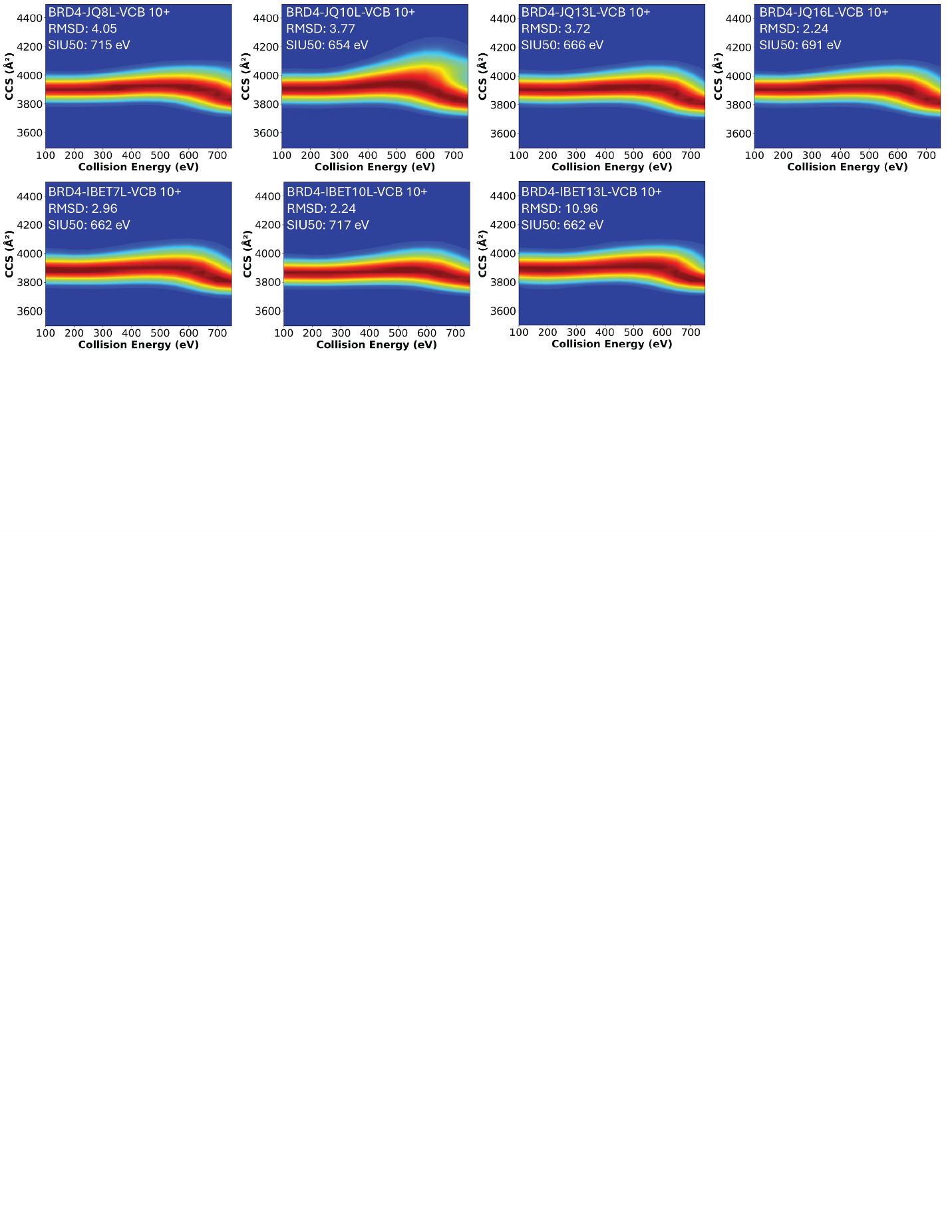

**Figure S9.** SIU plots for the 10+ charge state of each PROTAC complex studied with the JQ1 warhead (top row) and the I-BET726 warhead (bottom row) where linker length increases from left to right.
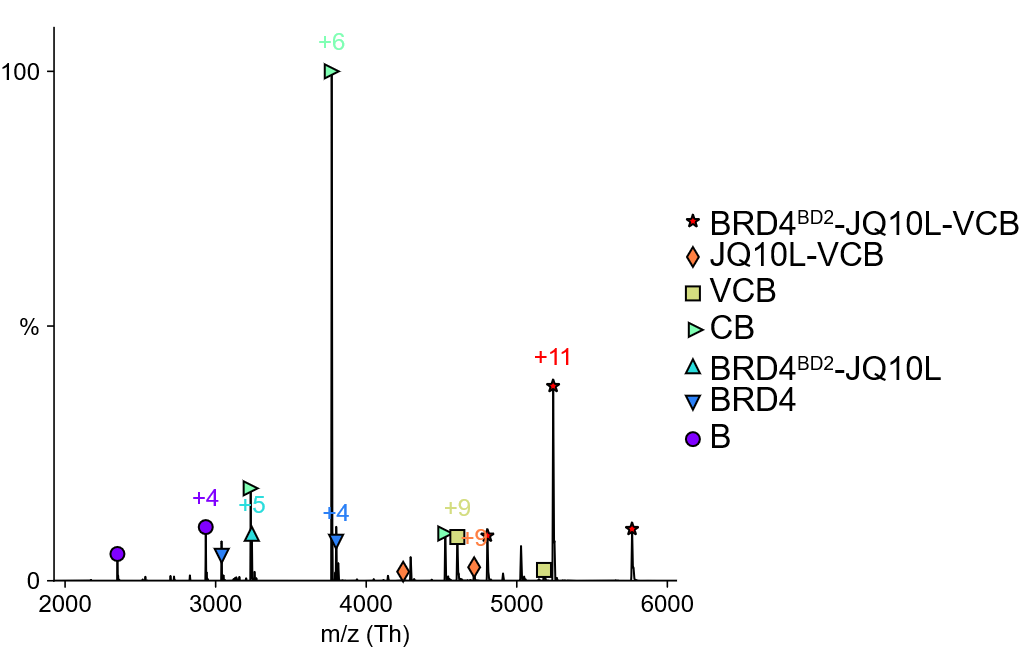

**Figure S10.** vT-ESI spectrum of BRD4^BD2^-JQ10L-VCB at 50 °C. A charge state has been assigned for each species present.

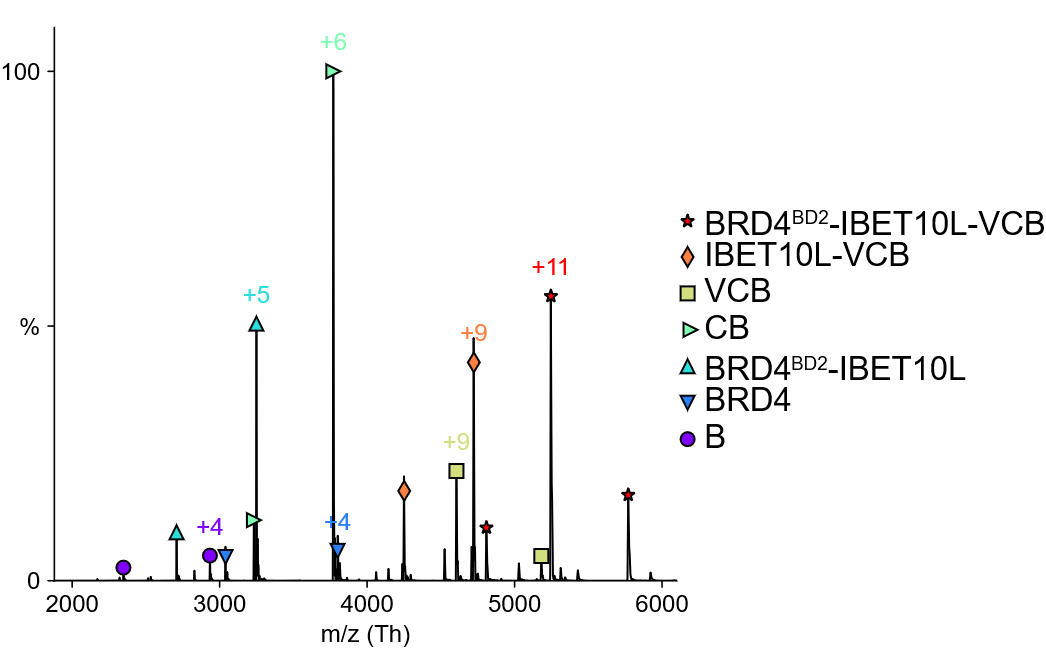

**Figure S11.** vT-ESI spectrum of BRD4^BD2^-IBET10L-VCB at 45 °C. A charge state has been assigned for each species present.

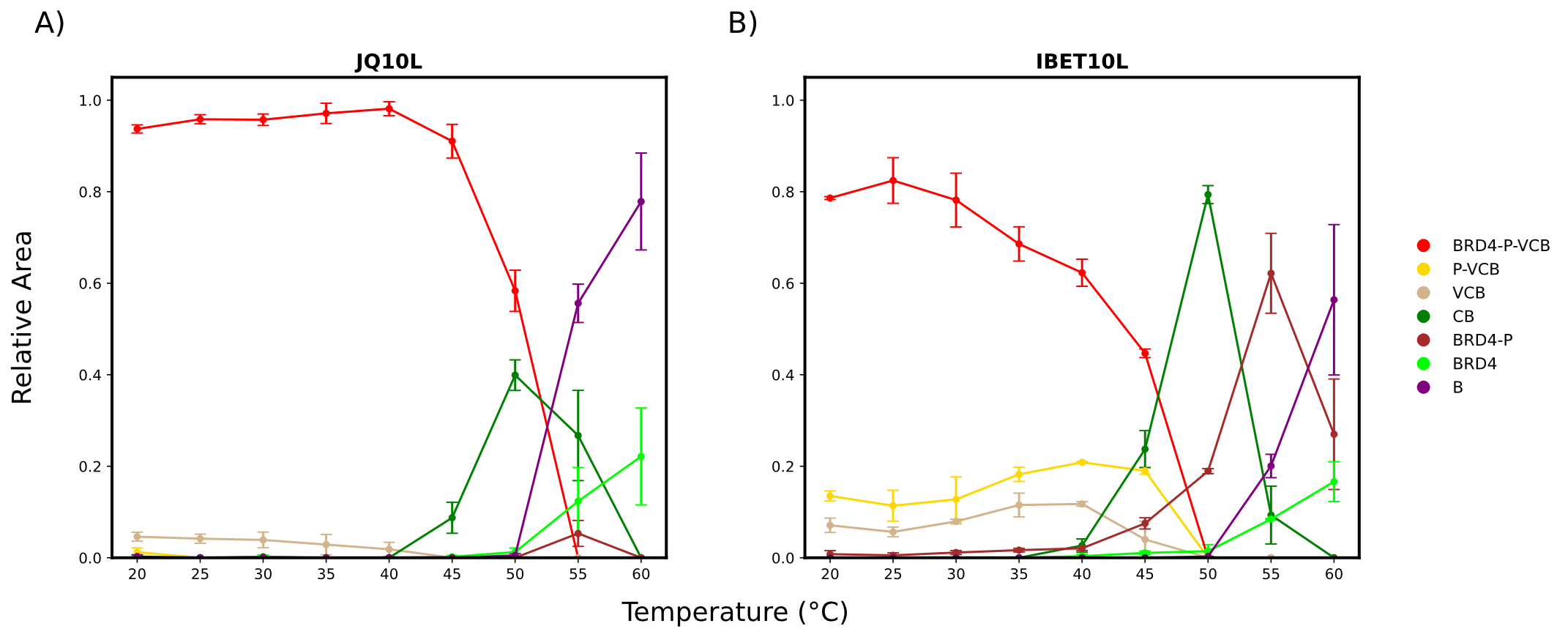

**Figure S12.** vT-ERMS plots for BRD4-JQ10L-VCB (A). vT-ERMS plot for BRD4^BD2^-IBET10L-VCB (B).

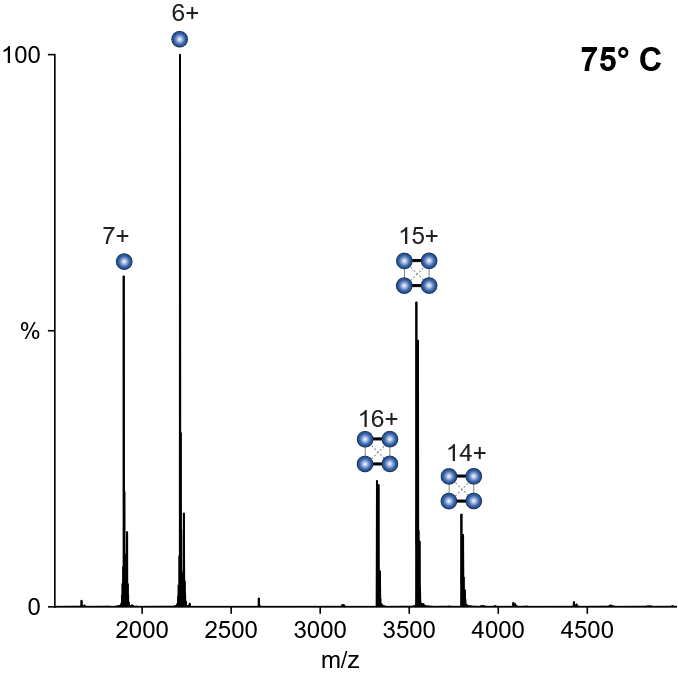

**Figure S13.** vT-ESI spectrum of streptavidin at 75 °C. Tetramer dissociates to monomer. In SID, tetramer dissociates to dimer, consistent with the known dimer-of-dimers structure of streptavidin.

| Lens | Transmission | SID 45 (V) | SID 45 + x (V) |
| --- | --- | --- | --- |
| Bent flatapole | 5 | 5 | 5 |
| Entrance 1-3 | -2 | -5 | -5 |
| Entrance 2 | -20 | 2 | 2 |
| Front top | 0 | -23.5 | -23.5+ 0.05x |
| Front bottom | 0 | -9.5 | -9.5 + 0.05x |
| Surface | 0 | -40 | -40 -x |
| Middle bottom | 0 | -110 | -110 -x |
| Back top | 0 | -154 | -154 -x |
| Back bottom | 0 | -90 | -90 -x |
| Exit 1-3 | -5 | -50 | -50 -x |
| Exit 2 | -22 | -30 | -30 -x |
| C-trap entrance inject | 1.8 | -29 | -29 -x |
| C-trap exit inject | -16 | -20 | -20 -x |
| C-trap offset | 0 | -35 | -35 -x |

**Table S3.** Voltages used for SID experiments on a modified Q Exactive Ultra High Mass Range (UHMR) Orbitrap Mass Spectrometer. SID voltage is defined by the bias between the bent flatapole and surface in the SID device.

| Sequence | Time | Pre Array Gradient | Pre Array Bias | Array Entrance | Wave Height | Array Offset | Array Mode | Array Exit | Post Array Gradient | Post Array Bias |
| --- | --- | --- | --- | --- | --- | --- | --- | --- | --- | --- |
| Inject | 10 | 85 | 74 | 10 | 6 | 60 | Forward | 50 | 35 | 10 |
| Separate | 5 | 85 | 70 | 30 | 0 | 70 | Sideways | 30 | 35 | 10 |
| Eject and Acquire | 144 | 85 | 70 | 50 | 25 | 45 | Forward Eject | 2 | 35 | 10 |

**Table S4.** Cyclic Control Ion Mobility Global Settings. Typical parameters used in cyclic sequence.
